# Widespread immunogenic poly-epitope frameshift mutations in microsatellite unstable tumors

**DOI:** 10.1101/662262

**Authors:** Vladimir Roudko, Cansu Cimen Bozkus, Theofano Orfanelli, Stephanie V. Blank, Benjamin Greenbaum, Nina Bhardwaj

## Abstract

Microsatellite instability-high (MSI-H) tumors are an important model system for evaluating neoantigen-based immunotherapies given their high tumor mutation burden and response to checkpoint blockade. We identified tumor-specific, frameshift peptides, encoding multiple epitopes that originated from indel mutations shared among patients with MSI-H endometrial, colorectal and stomach cancers. Epitopes derived from these shared frameshifts have high population occurrence rates, wide presence in many tumor subclones and are predicted to bind to the most frequent HLA alleles in the TCGA MSI-H patient cohorts. Neoantigens arising from these mutations are more dissimilar to both self and viral antigens, indicating the creation of peptides, that, when translated, can present truly novel antigens to the immune system. Finally, we validated the immunogenicity of common frameshift peptides from MSI-H endometrial patients in an array of T cell stimulation experiments, using peripheral blood mononuclear cells isolated from healthy donors. Our study describes for the first time the widespread occurrence and strong immunogenicity of tumor-specific antigens, derived from shared frameshift mutations in MSI-H cancer and Lynch syndrome patients, suitable for the design of common preventive “off-the-shelf” cancer vaccines.

## Introduction

Genetic alterations in tumor genomes that encode novel stretches of amino acids compared to normal cells are a potential source of immunogenic tumor-specific epitopes, commonly referred to neoantigens. Total neoantigen burden, (the sum of neoantigens predicted to be expressed by a tumor), has been demonstrated to be an independent proxy for response to anti-programmed cell death protein 1 (PD-1) immunotherapy^1–4^. However, determining neoepitopes in individual tumor sample remains fraught with uncertainties, such as the lack of congruence between selection of somatic variants and immunogenic neoantigens. Due to the presence of high loads of tumor-specific antigens, and strong effector T cell infiltration, microsatellite instability-high (MSI-H) tumors are an important model system for evaluating neoantigen-based immunotherapy in therapeutic and protective settings. The MSI-H tumor phenotype arises from defective DNA repair mechanisms due to a loss of the mismatch repair (MMR) activity. MSI-H is typically characterized by the variation of DNA length in microsatellite loci – units of one to ten mono-, di-, tri-, or tetra-nucleotides repeated multiple times^5^. In healthy cells these unpaired nucleotides are recognized and excised by MMR, but in MSI-H tumors they remain unrepaired. Some of these microsatellite regions are located in coding regions, where their destabilization may cause frameshift (fs-) mutations and thereby create a large somatic mutation burden, possessing a huge source of tumor-specific neoantigens.^4, 5^.

Inactivation of several MMR genes plays a key role in the acquisition of the MSI-H phenotype in hypermutated tumors^6–9^. This often happens at the later stages in tumorigenesis, and it originates by either deleterious somatic mutagenesis or epigenetic inactivation of MMR genes. This sporadic type of MSI-H tumors occurs in 10-40% of colorectal and endometrial cancers and is mainly caused by biallelic hypermethylation of the MLH1 promoter^14, 15^. Apart from somatic inactivation of MMR genes during tumorigenesis, germline mutations within the same genes are also found. Lynch syndrome, sometimes referred to hereditary nonpolyposis colorectal cancer (HNCC), is an inherited, autosomal-dominant disorder characterized by germline non-synonymous mutations in MRR genes, accounts for 3-5% of all colorectal and endometrial cancers. The majority of patients have germline mutations in the MSH2 (∼30%) and PMS2 (∼70%) genes^12^. Subjects with Lynch syndrome suffer from a high predisposition to develop cancer early life, with an 80% life time risk for colorectal or endometrial MSI-H cancers. Also estimates suggest as many as 1 in every 300 people may carry Lynch syndrome-associated germline alterations^13–15^. Epithelial tissues are primarily at risk of tumorigenesis, with colorectal cancers being the most common (80% of HNCC patients). The second most common cancer with a significant percentage of sporadic and hereditary-predisposed MSI-H type is endometrial carcinoma (60% of HNCC patients). The MSI-H group accounts for up to 28.6% of low-grade and 54.3% of high-grade endometrioid cancers. Other cancers such as bladder, gastric, ovarian, small bowel and renal are also somewhat predisposed^16^.

Most neoantigens that are predicted from point mutations are derived from patient-specific passenger mutations, with most shared driver mutations creating rarely presented peptides^17^. Consequently, the former are preferentially applied in cancer vaccine platforms. However, the MSI-H phenotype has several features which can be leveraged for common “off the shelf” vaccine design: (1) a high-mutational burden in well-defined, limited sequence space – microsatellite regions; (2) a restricted pattern of the mutations due to nucleotide insertions or deletions; and (3) a high probability of shared indel mutations in protein coding genes, which may induce formation of frameshift (fs-) peptide, which encodes multiple MHC-I restricted epitopes (poly-epitope fs-peptide) that are shared among multiple patients^18, 19^. Based on this premise, we investigated fs-mutations in tumor genomes of MSI-H patients with colorectal, stomach and endometrial carcinomas and identified a high frequency of broadly shared, immunogenic, poly-epitope fs-peptides.

## Results

### MSI-H colorectal, stomach and endometrial patient cohorts have a high fs-load

Though tumor evolution is primarily viewed as driven by a random mutational process, there is accumulated evidence that some mutations are acquired non-randomly^40–43^. Considering the existing skewness in mutational process and high load of indel mutations in microsatellite regions within open reading frames of MSI-H tumors^23^, we hypothesized MSI-H patients may share mutational events. While missense somatic mutations share limited similarity across multiple tumors^24, 25^, frameshift (fs-) mutations in microsatellite (MS) unstable regions are indeed likely to generate common fs-peptides (Supplement Figure 1A). Though previous studies have annotated somatic mutation load as well as the distribution and frequency of insertions/deletions on a pan-cancer scale^17, 18, 26^ (Supplement Figure 1B), little is known about the frequency of shared mutational events. We examined mutational load in cancer cell lines from Cancer Cell Line Encyclopedia^27, 28^ (CCLE) and found that fs-mutations are more frequenty shared among multiple cancer cell lines, than missense mutations (Supplement Figure 1C).

**Figure 1.**
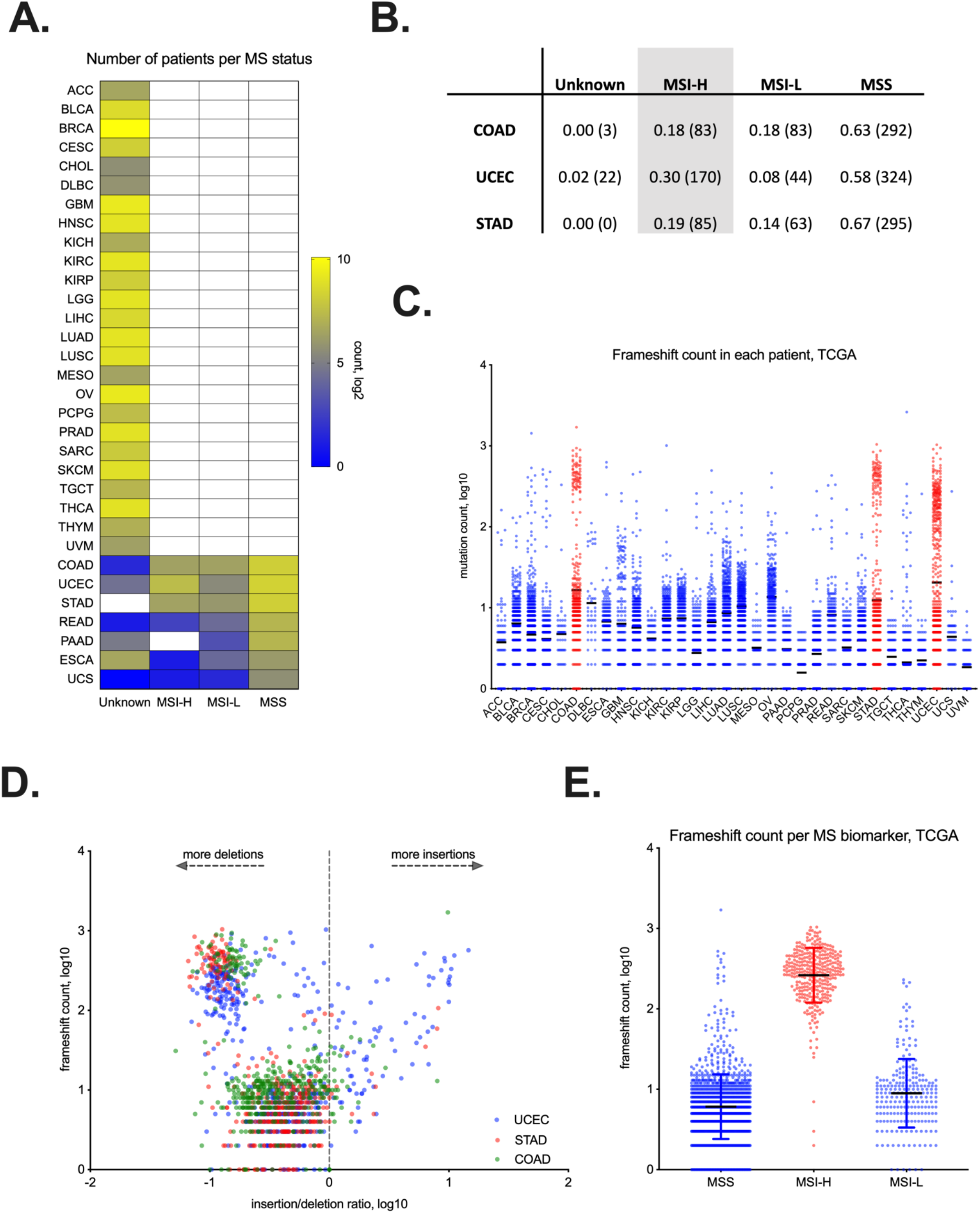
Microsatellite instability is detected in COAD, STAD and UCEC tumors in TCGA. Majority of MSI-H frameshifts are deletions. **A.** Quantification of patients with microsatellite instable (MSI) tumors according to biomarker, applied by TCGA. MSI-H - MSI-high, MSI-L - MSI-low, MSS - MS-stable, and Unknown – undetermined MS status. **B.** Table, showing the fraction (absolute number) of patients with UCEC, COAD and STAD tumors identified as MSI-H, MSI-L, MSS or Unknown. **C**. Frameshift (fs-) load (Y-axis, log10) in different tumor types across TCGA. **D.** Comparison of fs-load (Y-axis, log10) with insertion-deletion ratio (X-axis, log10) in COAD, STAD and UCEC tumors. **E**. Segregation of fs-load by MS biomarker.

The majority of TCGA tumors are microsatellite stable or their status is unknown. However, ∼20%-30% of endometrial, colorectal and stomach adenocarcinomas are diagnosed as microsatellite unstable, MSI-H (UCEC, COAD, STAD respectively, Figure 1A). In total, the MSI-H population in TCGA accounts for 338 patients (Figure 1B). The fs-load, as determined by fs-mutation count per each patient, is particularly high in a subset of UCEC, COAD and STAD patients (Figure 1C). Consistent with previous studies, the majority of frameshifts stems from nucleotide deletions, as determined by correlating patient’s fs-load with insertion-to-deletion ratio (Figure 1D). Finally, the majority of fs-enriched patients are segregated according to the clinical MSI-H biomarker, indicating nearly perfect specificity/selectivity of this biomarker in detecting indel-enriched tumor types (Figure 1E).

### Colorectal, stomach and endometrial MSI-H adenocarcinomas are enriched in shared poly-epitope fs-peptides

We then hypothesized that de-novo translated peptide segments – fs-peptides – within a newly-defined codon sequence frame (either +1 or +2) downstream of the indel mutation, may possess a significant source of immunogenic class I neoepitopes. If such fs-derived neoepitopes exist, they may also exhibit a high rate of turnover and processivity: mRNAs that encode frameshift mutations may be rapidly degraded through the nonsense-mediated decay (NMD) pathway, which is accompanied by nascent peptide decay on the 80S ribosome^29–33^. While the expression of fs-genes may be downregulated, the translated product is destabilized, potentially producing short, presentable peptides at a higher rate^34, 35^.

To identify potentially immunogenic epitopes derived from fs-mutations, we developed a fs-neoantigen calling pipeline (see **Online Methods**). Using this pipeline, we analyzed the distribution of fs-mutations, fs-peptides and corresponding fs-epitopes in MSI-H UCEC, COAD and STAD cohorts of TCGA patients. As expected, we found that many genes were commonly mutated via indels in MS regions in all the three tumor types. Up to 80% of MSI-H COAD and STAD patients had several commonly mutated genes as a result of indels, e.g. ACVR2A, MIKI67, RPL22. More than 50% of MS-H UCEC patients shared a different set of genes affected by frameshifts e.g. CASP5, MUC6, KMT2C. Expectedly, the frequency of shared fs-peptides, derived from exactly the same fs-mutations, dropped, with only a few fs-peptides shared by 40% of MSI-H UCEC, or 50% of MSI-H COAD/STAD patients. Finally, the frequency of shared HLA epitopes in MSI-H UCEC patients approached 25% having 28 immunogenic fs-epitopes, and to 50% of COAD patients who had 29 shared epitopes (Figure 2A, B). Interestingly, colon and stomach MSI-H tumors had twice as many shared immunogenic fs-events than endometrial MSI-H tumors (Figure 2B), which likely stems from the different pathways of tumorigenesis, and from the profiles of mutated oncogene drivers.

**Figure 2.**
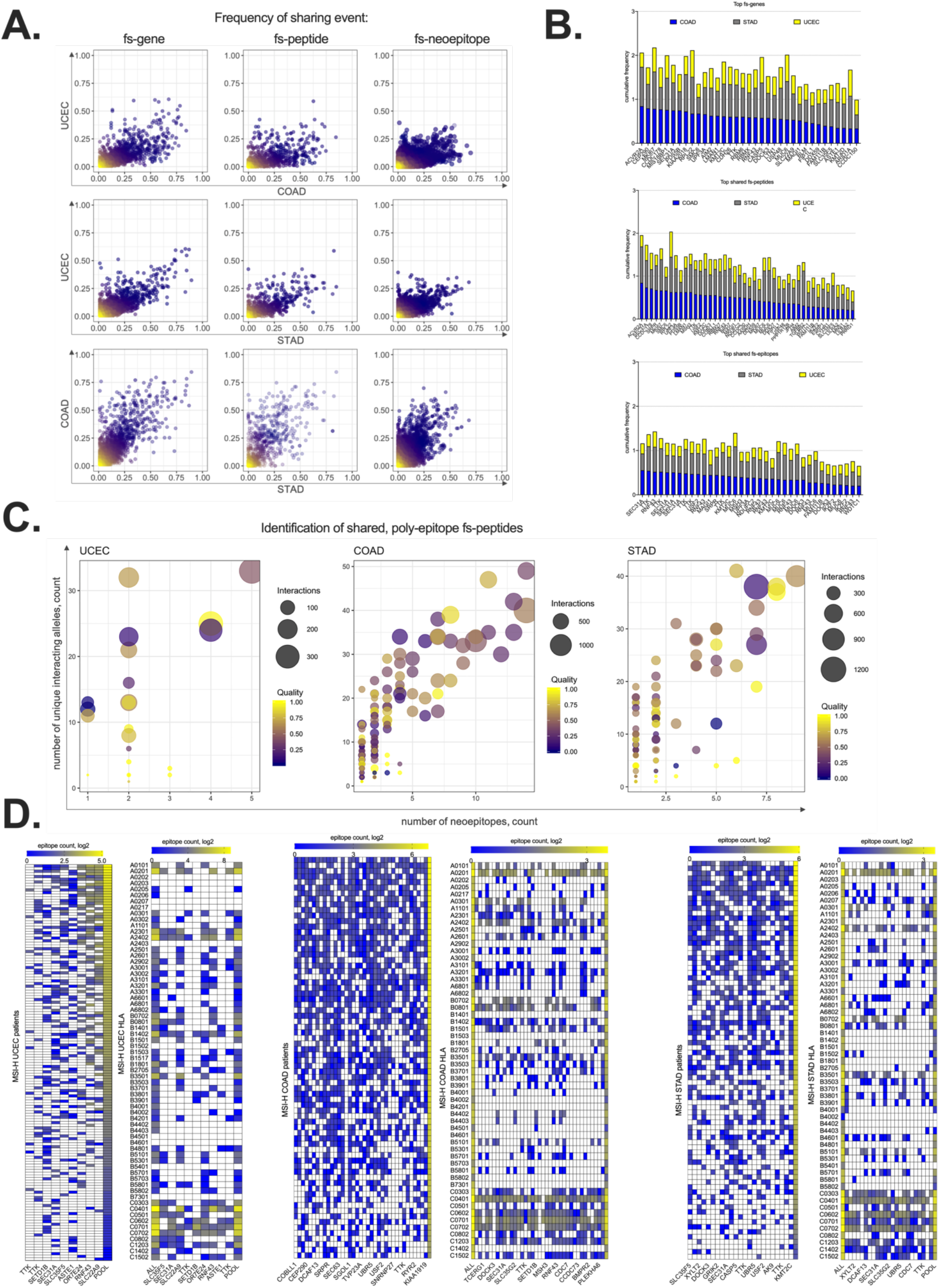
Frequencies of shared, fs-events in STAD, COAD and UCEC MSI-H tumors. Greedy pipeline for identification of shared, fs-peptides in MSI-H tumors. **A.** Scatterplots of patient frequencies of frameshifted genes (LEFT), fs-peptides (CENTER) and fs-epitopes (RIGHT) in UCEC, STAD and COAD MSI-H tumors. **B.** Cumulative frequency histograms, showing most frequently mutated genes via indels (TOP); genes, mutated by shared fs-peptide (CENTER); genes, encoding shared fs-neoantigens (BOTTOM). **C.** Three scatterplots showing selection procedure for identification of shared fs-peptides in UCEC (LEFT), COAD (CENTER) and STAD (RIGHT) patients. Each dot represents fs-peptide with the frequency of sharing above 20% in the patient cohort. Amount of predicted 9-mer epitopes per peptide (X-axis) is plotted against a number of predicted interacting HLA alleles (Y-axis). Size of the circle represents total number of predicted interactions between HLA-alleles and encoded epitopes. Color of the dot reflects the ratio of “PASS”ed fs-mutations to total number of called fs-mutations, according to somatic callers used by TCGA consortium. **D.** Quantification of T cell epitopes (log2 scale) derived from shared fs-peptides (columns) per each patient (rows, odd heatmaps) or each HLA allele (rows, even heatmaps) found in UCEC, COAD and STAD MSI-H cohorts.

To identify immunogenic fs-peptides with confidence, we developed a mutation ranking system based on maximization of four parameters (Figure 2C). Firstly, we introduced a quality metric for each called fs-mutation in which higher quality implies higher confidence that this mutation is truly somatic (Supplement Figure 2A). Also, we analyzed the distribution fs-peptide length: on average, the length of MSI-H frameshift peptides is 20-30 aminoacid (aa) residues, suggesting that these peptides may encode multiple immunogenic epitopes per fs-mutation (Supplement Figure 2B). Secondly, we maximized the number of putative neoepitopes per each fs-peptide: epitopes, predicted for each fs-peptide across all patients, were pooled and the total number of unique epitopes was determined. Thirdly, we grouped all HLA alleles predicted to bind those neoepitopes and considered this to be an HLA-ligandome of each fs-peptide. Maximization of this parameter allowed us to pick poly-allelic fs-peptides, covering as diverse a set of HLA alleles in a population as possible. Finally, to include the population HLA allele frequency parameter, we quantified total amount of antigen-allele interactions per frameshift. Together, we ensured the prediction of fs-peptides that are most likely immunogenic, encode poly-epitopes, bind to a broad spectrum of HLA-alleles and widely shared. We identified 9, 37 and 23 fs-peptides derived from frequently shared indel mutations that encode poly-epitopes in endometrial, colorectal and stomach MSI-H patients, respectively (Figure 2C). Counting the number of epitopes derived from each fs-peptide in each patient showed a high level of presence of these epitopes in MSI-H cohort of patients. Alternatively, plotting the number of epitopes derived from frameshift sequence per each predicted HLA allele showed that many epitopes are predicted to bind to the most frequently expressed HLA alleles in the TCGA patient cohort (Figure 2D). Finally, among the detected fs-peptides, five were found in all MSI-H patients, pointing to the possibility to develop an off-the-shelf MSI-H vaccine for these three tumor types: SLC35F5, SEC31A, TTK, SETD1B and RNF43.

### Nine poly-epitope fs-peptides shared in MSI-H endometrial carcinoma originate from clonal tumor-specific alleles and are correctly translated

We next focused our attention on frameshift-derived neoantigens from MSI-H endometrial carcinoma, as these have not been characterized to date. At least two neoepitopes from nine selected fs-peptides are detected in >95% of the MSI-H UCEC TCGA patient cohort. Moreover, binding profiles of the predicted epitopes cover almost all HLA-alleles of the same patient population as well as targeting the majority of frequent HLA types (Figure 2D). Importantly, only the “mixture” of MHC-I epitopes derived from all 9 peptides has the potential to reach a significantly high representation of all HLA allele-epitope diversity, than each fs-peptide separately (Figure 3A).

**Figure 3.**
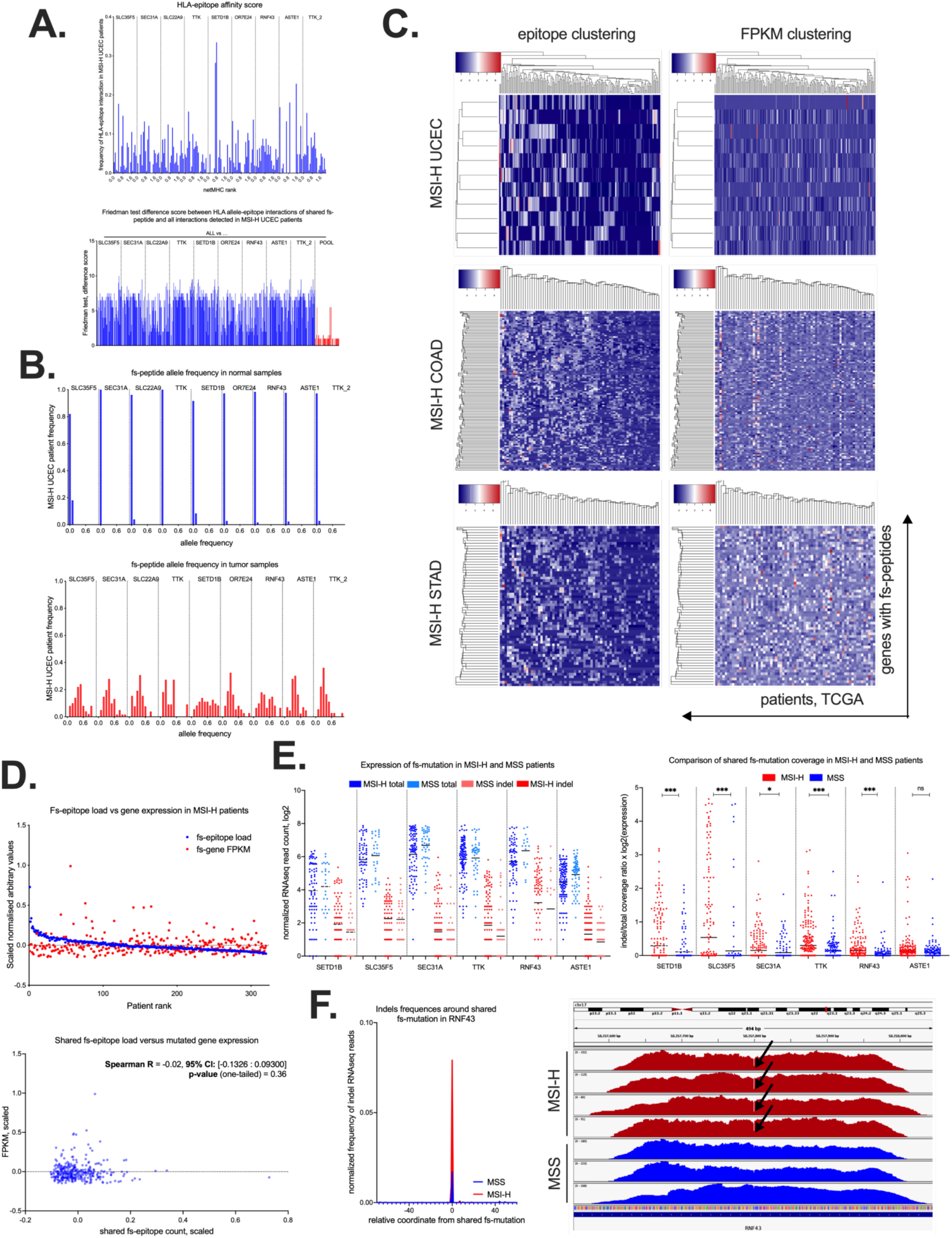
Genomic and expression properties of shared, immunogenic fs-mutations in MSI-H UCEC tumor. Neoantigen quality of predicted epitopes. **A.** TOP – Distribution of predicted HLA allele-epitope interaction ranking scores (NetMHC v3.4/v4.0), derived from nine shared fs-peptides in MSI-H UCEC. BOTTOM – Friedman difference score, measuring the difference between total amount of HLA-allele epitope interactions found in MSI-H UCEC patients (ALL) and either sum of epitope-allele interactions per each fs-peptide separately, or pooled together (POOL). Each bar represents the difference score between number of predicted epitopes per each HLA allele. **B.** tumor allele frequency of nine shared frameshift mutations in normal (TOP) and tumor (BOTTOM) tissues of MSI-H UCEC TCGA patients. **C.** Unsupervised hierarchical clustering of shared fs-mutations (LEFT) and corresponding FPKM values of frameshifted genes (RIGHT) in MSI-H UCEC (TOP), COAD (CENTER) and STAD (BOTTOM) tumors. Patients plotted in columns, genes plotted in rows. **D.** TOP – scatterplot of shared fs-load and corresponding gene expression per each MSI-H patient. BOTTOM – correlation plot between scaled fs-load and expression. **E**. Normalized expression of nine shared fs-mutations in MSI-H UCEC patients. LEFT – normalized expression of total and indel-containing reads spanning the genomic loci of six shared fs-mutations. RIGHT – ratio of indel to total read count spanning the microsatellite region in MSI-H and MSS patient cohort of UCEC, STAD and COAD tumors. Statistical significance is derived from non-parametric Mann-Whitney two-tailed test. **F.** Representative example of shared frameshift detected in RNAseq datasets. LEFT – normalized frequency of fs-mutation in RNF43 within 100 nucleotide genomic loci: 50 nt upstream and 50 nt downstream the shared frameshift in MSI-H and MSS RNAseq samples. RIGHT – RNAseq read histograms around the shared frameshift in RNF43 from MSI-H (red) and MSS (blue) patients. The 1-nucleotide long drop in RNAseq read coverage at the site of shared frameshift is indicated with an arrow.

Additionally, we analyzed the tumor allele frequencies from the MSI-H UCEC patient cohort to estimate the abundance of the selected nine shared frameshifts in tumor genomes. To do that, we compared corresponding fs-allele frequencies in normal and tumor samples. As expected, fs-alleles are barely detectable in normal tissues, but in tumor biopsies their frequencies rise up to 40% on average in tumor biopsies, indicating these mutations are often clonal (Figure 3B). This suggests the 9-peptide mix has a potential to prime T cell responses against almost all malignant cells. However, the high mutation rates of MSI-H tumors might decrease the probability of the shared fs-peptides being translated (Supplementary Figure 2C). Taking this into consideration, we assessed the conditional probability of shared fs-peptide being correctly translated given this shared mutation has happened. For this purpose, we estimated all disruptive upstream and downstream mutation frequencies using the MSI-H TCGA patients and calculated posterior probabilities (Supplementary Figure 2D). Though the majority of MSI-H UCEC shared fs-peptides has a high probability of being correct (>0.9), TTK and RNF43 frameshifts have an increased chance of being inactivated, with a ∼0.8 probability of being correct.

### The predicted nine shared poly-epitope fs-peptides are expressed in MSI-H UCEC patients

To estimate the expression level of genes encoding shared fs-peptides, we analyzed RNA expression derived from TCGA RNAseq samples of matched MSI-H patients (Figure 3C-F). We performed unsupervised clustering of MSI-H patients by genes with predicted shared fs-events, obtaining the patient and gene rankings according to the shared fs-load (Figure 3C). We then plotted FPKM expression values of genes encoding shared fs-peptides, and ranked them according to the previously obtained patient and gene rankings (Figure 3C). We observed no correlation between RNA expression and shared fs-load, suggesting that frameshifted genes are not selectively epigenetically silenced in tumors (Figure 3D). We next analyzed the expression of nine shared fs-mutations in MSI-H UCEC patients. As two of the shared fs-mutations are present in one gene, formally we detected 8 uniquely mutated genes, and six of these were expressed at the RNA level from matching tumor samples (Figure 3E). To assess the expression level of fs-alleles, we compared the normalized read count containing the indel with total amount of reads covering the targeted genomic loci from MSI-H RNAseq samples and MSS samples as a control (Figure 3E). We detected statistically significant and robust expression of fs-alleles in MSI-H patients (Figure 3E). Also, we showed representative RNAseq expression histograms, covering the predicted shared frameshift in RNF43 gene, and derived indel RNAseq read frequencies in MSI-H and MSS patients (Figure 3F). Thus, we confirmed that fs-alleles are indeed expressed and detected in bulk RNAseq.

We then tested if high load of shared poly-epitope fs-mutations had a survival benefit within the MSI-H patient cohort and/or may be skewed towards later tumor stage. This questions is important for ongoing immunotherapy trials, where MSI-H becomes widely accepted as a predictive biomarker to anti-PD-1 therapy^36, 37^, although deeper genetic differences between patients may underlie the incomplete responsiveness. For this purpose, we performed Cox regression and survival analyses of TCGA MSI-H UCEC patients stratified by shared fs-load, as high and low, and analyzed tumor stage and patient age in the same strata. Patient age and tumor stage were evenly represented in both, fs-high and fs-low MSI-H cohorts (Supplemental Figure 3). Finally, we did not detect any significant benefit in patients’ survival based on shared fs-load in any of the MSI-H tumor types (Supplementary Figure 4).

### Tumor fs-epitopes are more likely to be presented and are less similar to viral antigens than missense derived epitopes

We next examined the intrinsic properties of fs-derived epitopes when compared to missense-derived epitopes and viral antigens. In previously published studies, similarity of tumor-derived neoantigens to pathogen derived (viral) antigens has been shown to be predictive in the checkpoint blockade immunotherapy setting^1, 2^. First, we calculated the total amount of neoantigens derived from missense mutations of MSI-H patients and compared it to fs-epitope load. Despite the fact that the total frameshift and the missense mutation loads were similar, the number of MHC-I epitopes per mutation were different: 4 epitopes per one frameshift and 2 per one missense mutation on average (Supplement Figure 5A). This observation suggests, that fs-mutations may be more immunogenic then missense due to an increased probability of presentation. While most of missense-derived epitopes are, by definition, one aa different from a self-peptide, the majority of fs-derived epitopes are unique, “non-host” peptide sequences and hence exhibit minimal similarity to the human proteome (Supplement Figure 3B). This implies, that fs-derived epitopes might have never been tolerized by the host immune system, and that the frameshift-specific T-cells may have little or minimal autoreactivity.

We also compared these two epitope datasets with virus-derived antigens. At multiple search parameters, the overall number of missense epitopes, when matched with viral ones, was higher, than matched fs-epitopes (Supplement Figure 3C). We speculate this observation is due to the overall viral adaptation to the human proteome and host T-cell epitopes, as viruses need to hijack particular host functionalities in order to interact with the host cellular machinery as well as escape host immune recognition. As a consequence of this finding, we conclude fs-epitopes appear “less self’ than either missense or virus derived epitopes.

### Predicted shared poly-epitope fs-peptides are detected and expressed in diverse set of cancer cell lines from CCLE

To validate the predicted shared fs-mutations in an external dataset, we verified the presence of these mutations in the Cancer Cell Line Encyclopedia (CCLE) (Figure 4). 34 from 45 of predicted shared poly-epitope fs-peptides were detected in multiple cancer cell lines, derived from different tumor types. Diverse cancer cell lines had different numbers of shared fs-peptides, e.g. intestine, endometrium, stomach and prostate cancer cell lines having 5-10 shared fs-mutations per cell line, while hematopoietic, ovarian and lung cancer cell lines had 1-5 shared fs-mutations per cell line (Figure 4A). The presence of predicted shared fs-mutations in the last three tumor types suggests a broader occurrence of shared poly-epitope fs-peptides across tumors, than initially suggested. The frequency of shared fs-positive cell lines per tissue type was around 20, 40 and 60% of intestine, stomach and endometrial cell lines (Figure 4B). Interestingly, the majority of fs-mutations differentially shared by MSI-H UCEC, COAD and STAD TCGA tumors were evenly detected in endometrial, intestine and stomach cancer cell lines (Figure 4C), suggesting epigenetic remodeling of certain chromatin areas making them accessible for genome destabilization. Initially predicted in TCGA, shared fs-peptides were the only fs-mutations statistically significantly shared in cancer cell lines than any other fs-mutation derived from the same genes (Figure 4D). Fs-allele coverage analysis suggested that predicted shared fs-mutations are clonal and present in shared fs-positive cancer cell lines at 30-50% allele frequency on average (Figure 4E). Also, RNAseq data indicates the gene expression patterns are unchanged upon acquiring shared frameshift (Supplement Figure 6). Overall, the CCLE dataset analysis confirms the widespread occurrence of predicted shared fs-peptides in cancer.

**Figure 4.**
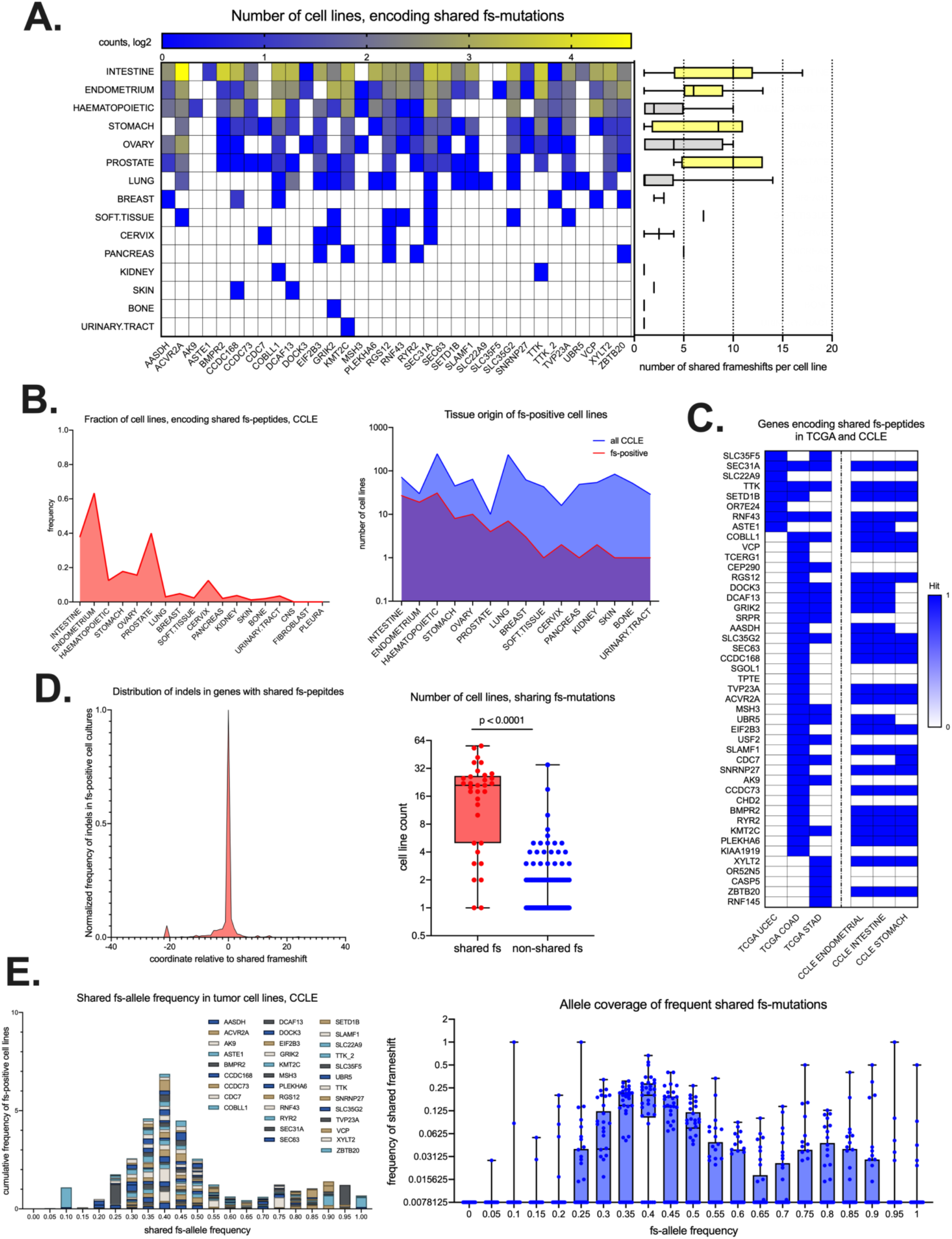
Detection of shared fs-mutations in cancer cell line encyclopedia (CCLE). **A.** Quantification of shared fs-mutations in cell lines per tissue of tumor origin and per each frameshifted gene. Histogram plot on the left shows number of shared fs-mutations per cell line. **B.** General statistics of cell lines, encoding shared fs-peptides. LEFT – fraction of cell lines, positive for shared fs-mutation, per each tissue type. RIGHT – Absolute number of cell lines with detected shared fs-mutation compared to total number of cell lines in CCLE. Cell lines are sorted according to tissue origin. **C.** Gene and tumor specificity of shared fs-mutations in TCGA and CCLE. **D.** Distribution of all detected indels in genes with shared fs-mutations (34 genes in CCLE). LEFT – metagene, showing normalized frequency of all detected indels in 34 genes, around shared fs-mutation. RIGHT – t-test of number of cell lines encoding shared fs-mutation versus all other fs-mutations, detected in selected 34 genes. **E.** fs-allele frequency in WES samples of shared fs-mutation positive cell lines. LEFT – cumulative histogram plot, showing fs-allele frequency of each shared fs-mutation per gene of origin. RIGHT – box-plot of fs-allele frequency plotted against frequency of fs-mutation in the pool of fs-positive cell lines.

### Nine poly-epitope fs-peptides shared in MSI-H UCEC patients are highly immunogenic

To assess the immunogenicity of the nine predicted fs-peptides from MSI-H UCEC patient cohort, we evaluated the T cell responses against each neopeptide using a T cell immunogenicity assay developed in our laboratory that is designed to rapidly prime naïve T cells (see **Online Methods**). In brief, long overlapping peptide (OLP) libraries spanning each fs-peptide were designed (Supplementary Figure 7). Using these OLP pools, T cells from 15 randomly picked healthy donors (HD) were primed and expanded. After expansion, the cells were stimulated with the OLP pools and fs-peptide-specific T cell responses were evaluated by measuring IFN-γ production using ELISPOT (Figure 5A). Results showed that each fs-peptide could elicit T cells responses in a subset of subjects tested. Furthermore, some subjects had reactive T cells against multiple fs-peptides (Figure 5B-D). Importantly, when combined, the fs-peptide-specific T cells were significantly enriched in the subject cohort in accordance with the predictions detailed in Figure 2D. We also characterized the fs-peptide-specific T cell responses in the same HD cohort by intracellular staining (ICS). Results from both assays showed similar stimulation profiles. Moreover, responses to fs-peptides were observed primarily in CD8+ T cells, reaching up to 10% of T cells indicating strong priming to these neoantigens. (Figure 5E-G). In total, a majority of HD (13 out of 15) responded to at least one fs-peptide. Importantly, the reactive T cells produced TNF-α, in addition to IFN-γ, suggesting that fs-peptide-specific T cells are polyfunctional (Supplementary Figure 8). Additionally, we synthesized control peptides (15-aa) for each fs-peptide using their wild type sequence surrounding the fs-mutation site. Responses by HD T cells to stimulation with WT OLP pool were not higher than the background (Figure 5H), suggesting that the observed T cell responses were specific to fs-peptides. Next, we investigated whether the fs-peptide-specific T cells responses that were observed in the HD cohort correlated with the predicted high affinity epitope load. To determine the predicted epitope load, we identified the HLA-I alleles of each subject by sequence-based HLA-I genotyping and investigated the predicted binding affinity of epitopes from fs-peptides to each subject’s unique HLA. We found no significant correlation between the total epitope load per patient and experimentally observed response rate (Figure 5I-K). This observation is expected since many studies have reported that the majority of predicted epitopes fail to elicit T cell responses^38^ and that high HLA-peptide binding affinity does not equate to immunogenicity. Altogether, our data show that MSI high patients have an increased frequency of high-quality T cell epitopes derived from shared fs-peptides, binding to a broad spectrum of HLA alleles, capable of inducing immunogenicity for CD8+ T cell in particular.

**Figure 5.**
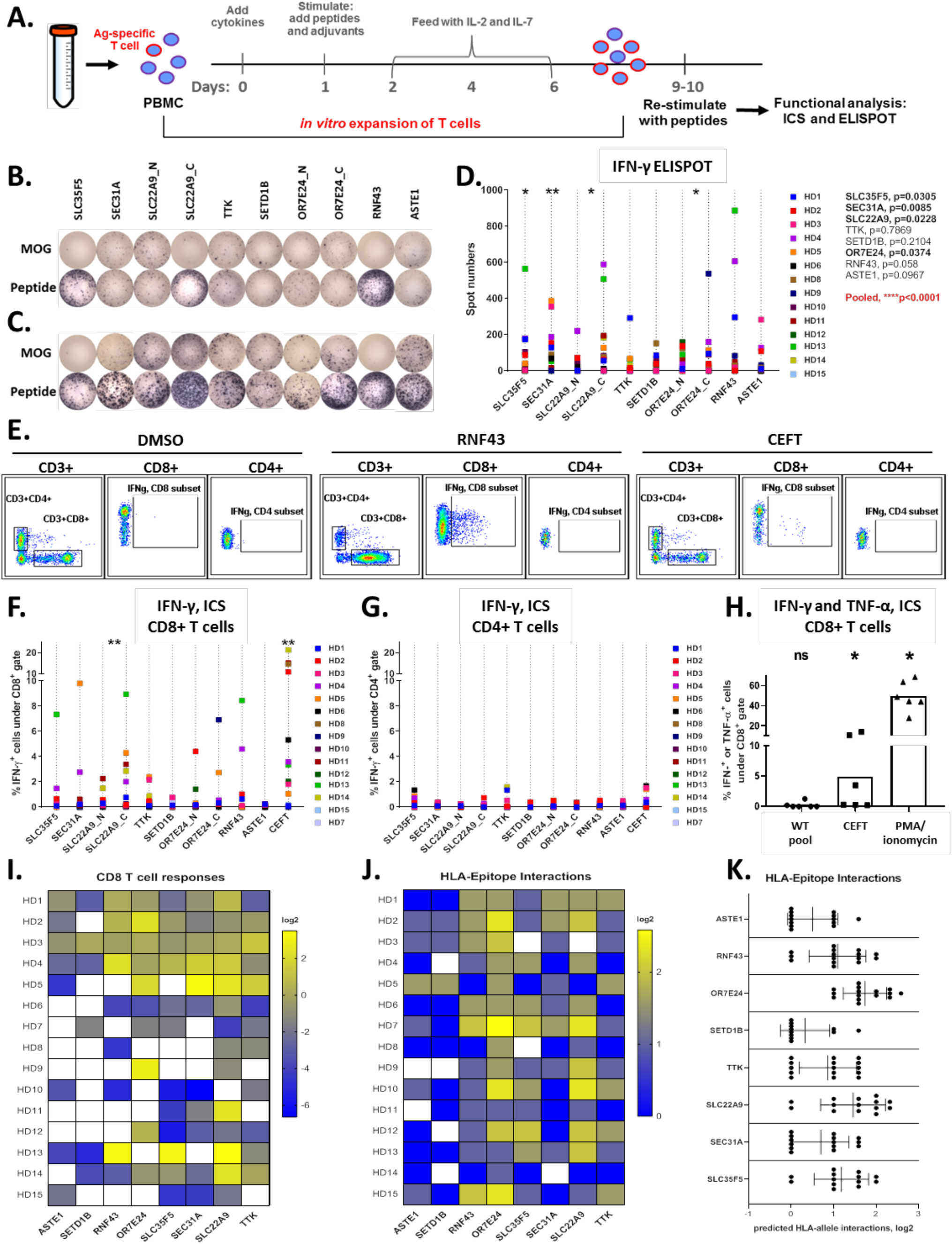
Shared fs-peptides predicted from UCEC MSI-H patients elicit T cell responses. **A.** An overview of T cell immunogenicity assay used to evaluate antigen-specific T cell responses. PBMCs from healthy donors (HD) were expanded *in vitro* following stimulation with fs-peptide OLPs as shown in supplementary figure 7. Expanded T cells (5×10^4^ cells/well) were re-stimulated with either the peptide pool they were expanded with or the control peptide pool MOG. Representative IFN-γ ELISPOT images for **B.** HD13 or **C.** for selected responsive HD. **D.** Summary of ELISPOT data (n=14). Statistical significance for MOG vs OLPs was evaluated by Wilcoxon signed-rank test. **E.** Representative flow cytometry plots and summary of data (n=15) for IFN-γ in **F.** CD8 and **G.** CD4 T cell subsets. Stimulation with CEFT was used as a control. Statistical significance for DMSO vs OLPs was evaluated by Wilcoxon signed-rank test. **p=0.0032 for SLC22A9 and **0.0031 for CEFT. **H.** Frequency of IFN-γ or TNF-α producing CD8+ T cells upon stimulation with WT OLP pool. CEFT and PMA/Ionomycin stimulation was used as a control. The spot numbers and % IFN-γ values were calculated by subtracting the values obtained after MOG or DMSO stimulation from the values after OLP pool stimulation and negative values were set to zero. **I.** Summary of log2 transformed data for IFN-*γ* /TNF-*α* response by CD8+ T cells against fs-peptides. **J.** and **K.** Quantification (log2 transformed) of HLA allele-epitope interactions for each subject per fs-peptide. Interactions were counted when IC50<500 or percentile rank<2.

## Discussion

In this study we evaluated the MSI-H patients from the TCGA database for the presence of shared, immunogenic tumor-associated neoantigens, providing a basis for the design of a common “off-the-shelf” cancer vaccine. We proposed fs-events to have a high probability of being shared across multiple MSI-H patients. To this end, we characterized the MSI-H population and the tumor genomic frequencies of corresponding indel mutations, as well as their tumor expression profiles. Our approach to detect immunogenic, shared neoantigens relies on two assumptions: (i) indel mutations occurring in frequently mutated MS regions will lead to identical fs-peptide extensions, (ii) these frequent, identical fs-peptide extensions encode poly-epitopes with broad HLA allele specificity. We confirmed the validity of our neoantigen selection approach by testing the immunogenicity of the selected fs-peptides. We found that the selected peptides were highly immunogenic and generated strong CD3+CD8+ T cell responses in a broad range of subjects.

One important finding is the existence of shared mutations that generate immunogenic neoantigens. Indeed, the majority of tumors develop a restricted profile of mutations, either disrupting activity of oncosuppressors, like p53 and KREB, or promoting gain-of-function activity of oncogenes, like BRAF V600E. However, these shared mutations often escape immune recognition and are less likely to encode tumor-associated neoantigens^17^. By contrast, in MSI-H tumors indel mutagenesis is generally restricted to MS regions, thus increasing the probability of being shared among patients (Supplement Figure 1A, B). Another key observation is that indel mutations in MS within protein coding regions are more likely to produce common fs-peptide, which may encode immunogenic neoantigens. With an average length of 15 – 50 aa residues, frameshift extensions can easily accommodate multiple 9-mer T cell epitopes. By running conventional antigen-prediction pipelines, we annotated several shared fs-peptides as immunogenic.

From the tumor’s evolutionary perspective, there is no need to keep those immunogenic mutations, so multiple tumor-intrinsic escape mechanisms might exist as a consequence. As described elsewhere, MSI-H tumors upregulate checkpoint molecules to evade the development of antitumor T cell responses^12, 26, 37, 39^. Blockade of this mechanism has been proven to be effective in clearing a range of MSI-H tumors in multiple clinical trials^37, 40^. However, several other immune resistance mechanisms might exist: downregulation of HLA-allele and/or *β*-microglobulin expression; inactivation/loss of antigen processing genes; inactivation/loss of interferon-*γ*-response pathway genes; disruption of immunogenic neoantigens by additional, acquired mutations^41–43^. The latter have been examined by us with respect to mutational escape of shared poly-epitope fs-peptides. Indeed, in certain cases we observed mutations that were potentially disruptive to the predicted neoantigens (Supplementary Figure 2). Another important finding is the high-occurrence of predicted shared fs-peptides in commonly utilized cancer cell lines (CCLE, Figure 4). We therefore suggest the potential utility of these shared fs-positive cell lines in immunological studies of predicted fs-epitopes, such as understanding the efficacy of antigen-dependent tumor cell killing as well as studying prospective tumor-escape mechanisms.

Finally, we investigated the sequence space produced by fs-mutations. Previous reports provided evidence that high quality immunogenic neoantigens are related to viral epitope sequences and are predictive for outcome of checkpoint blockade immunotherapy^1, 2^. To determine whether the same rules applied to fs-derived neoantigens, we investigated similarities between viral epitopes and immunogenic neoantigens. We found missense-derived neoantigens to be 3 times more similar to viral epitopes than fs-neoantigens. We attributed this to viruses evolving within the cellular environment of their host, trying to mimic the host’s functionalities and potentially avoid immune responses through extensive similarity to the host proteome. The fs-mutations are therefore even “further from self” than viral antigens (Supplemental Figure 3). Taken together, we conclude that frameshifts represent unique, intrinsically different epitope space, with a great potential for discovering immunogenic epitopes which can be targeted by immune therapies. In a recently published study authors investigated the presence of fs-derived neoepitopes in TCGA and arrived at similar conclusions^44^. Also, a few previous reports investigated the immunogenicity of unique fs-mutations on a small scale ^45-49^. In addition to this shared research goal, our group performed in depth computational characterization and experimental validation of shared fs-epitopes in MSI-H tumors specifically.

Preselected fs-mutations might also be used for developing targeted sequencing panels for diagnostic purposes. Precise detection and mapping of predicted mutations in each tumor patient may enhance the chances of achieving a positive response with an applied vaccine. The usage of targeted sequencing panels for diagnostics have already proven essential for developing actionable treatments, particularly in the selection of targeted regimens. We believe the same paradigm will become useful for precise immunotherapies, with physicians being able to select the ideal cancer vaccine formulation based on the results of targeted sequencing panels.

Our work revealed the possibility of designing common cancer vaccines in specific tumor subtypes with broad HLA-allele specificity. By applying tailored vaccines for MSI-H endometrial, colorectal and stomach carcinomas, one can potentially achieve immunological responses against existing neoplasms or develop preventive memory T cell responses in high-risk patient populations, like those with Lynch syndrome.

## Online Methods

### Computational analysis of mutational data from TCGA

Tumor-associated antigens were predicted using somatic mutation datasets, called by internal mutation pipelines of The Cancer Genome Atlas (TCGA, or Genomics Data Commons, https://gdc.cancer.gov). Briefly, annotated somatic missense and frameshift mutations by Mutect, Somatic Sniper, Varscan and Muse were combined together per each patient. In case of somatic missense mutations, corresponding 17-aminoacid residue-length normal peptides, surrounding mutation site, were converted to tumor-specific peptides and used for MHC-I epitope prediction. In case of frameshift mutations, the tumor specific peptide was called as following: major mRNA isoform was mutated according to the frameshift mutation, translated starting from “-8” aminoacid residue position from the mutation site till the stop codon within the new open reading frame, defined by the frameshift. Resulting frameshift peptides were used for MHC-I epitope prediction. NetMHC v4.0 and NetMHCpan v3.0^50, 51^ were used to predict missense and frameshift epitopes. HLA allele types for >5000 patients from TCGA were taken from previously published paper^3^. Collected epitope data was analyzed using statistical packages, available in Prism and R, using custom written scripts.

### Expression analysis of TCGA

Hg19-aligned RNAseq bam files were downloaded from GDC (https://gdc.cancer.gov). Obtained .bam files were processed with samtools to extract RNAseq reads, covering 250 nt genomic loci around shared fs-mutation (samtools view -b -L chr19.region.bed $i/*.bam > $filename.chr19.bam). To count indel events in extracted RNAseq bam fiels, we applied samtools mpileup tool: samtools mpileup -uf /work/scratch/index/TCGA/GRCh38.d1.vd1.fa $i | bcftools view -l region.bed - | grep “INDEL” >> $filename.vcf. Finally, obtained data was processed with custom scripts and analysed in PRISM 8.

### Peptide comparison with virus epitope databases

The collection of viral MHC-I epitopes was downloaded from IEDB database and preformatted for BLAST usage (makeblastdb -in iedb.fasta -parse_seqids -dbtype prot). Predicted frameshift and missense originated T-cell epitopes form MSI-H patients were BLASTed against IEDB. For comparison with viral we used following command: blastp -db iedb.fasta -query frameshift.neoantigen.netMHC.score.fasta -outfmt “6 qseqid sseqid pident ppos positive mismatch gapopen length qlen slen qstart qend sstart send qseq sseq evalue bitscore” -word_size 3 -gapopen 32767 -gapextend 32767 -evalue 1 -max_hsps_per_subject 1 -matrix BLOSUM62 - max_target_seqs 10000000 -out frameshift.neoantigen.iedb.blast.out. To compare predicted epitopes with human proteome, we used last command. First, we preformatted human proteome (ensemble archive from December 2016): lastdb -p human.proteome human.proteome.fasta. Then we used following command to compare epitopes to this database: lastal -f MAF -r 2 -q 1 - m 100000000 -a 100000 -d 15 -l 4 -k 1 -j1 -P 10 human.proteome frameshift.neoantigen.netMHC.score.fasta > frameshift.neoantigen.human.last.out. Finally, obtained results were processed with custom scripts (bash, python) and analyzed in PRISM 8.

### Computational analysis of CCLE database

Somatic mutation data and normalized RNAseq expression values for genes with shared fs-mutations were obtained from https://portals.broadinstitute.org/ccle. Total mutation counts per all cell lines were taken from CCLE_DepMap_18q3_maf_20180718.txt. Obtained data was processed with custom scripts and analyzed in PRISM 8.

### Code availability

The scripts and code are freely available on git account: https://github.com/VladimirRoudko

### Peptide synthesis

Custom peptide libraries for WT and mutated peptides were chemically synthesized by GenScript (USA/China). Each peptide had >80% purity as determined by high performance liquid chromatography. MOG and CEFT peptide pools were commercially available at JPT Peptide Technologies (Germany). Each peptide was resuspended in DMSO and used at a final concentration of 1 µg/mL. Sequences of mutated peptides are displayed in supplemental figure 7 and the WT sequences are as follows: for SLC35F5 GKLTATQVAKISFFF, for SEC31A QAVQSQGFINYCQKK, for SLC22A9 LEILKSTMKKELEAA, for TTK ESHNSSSSKTFEKKR and YSGGESHNSSSSKTF, for SETD1B MENSHPPHHHHQQPP, for OR7E24 MSYFPILFFFFLKRC, for RNF43 KSSLSARHPQRKRRG and for ASTE1 AEIFLPKGRSNSKKK.

### Rapid T-cell activation protocol

Healthy donor PBMCs were cultured in X-VIVO15 media (LONZA) with cytokines promoting dendritic cell (DC) differentiation overnight and then stimulated with peptide pool as displayed in supplemental figure 7 (each peptide at 1 μg/mL) in the presence of adjuvant promoting DC maturation in X-VIVO15. Stimulation with DMSO (vehicle) and MOG pool (JPT, 1 μg/mL) were used as negative controls and CEFT pool (JPT, 1 μg/mL) were used as positive controls. Next day, cells were fed with IL-2 (R&D Systems, 10 IU/mL) and IL-7 (R&D Systems, 10 ng/mL) in RPMI media (Gibco) containing 10% human serum. Cells were fed every 2-3 days. IL-2 and IL-7 were not added at the last feeding. After 10 days of culture, cells were harvested and re-stimulated with peptides (1 μg/mL) in the presence of anti-CD28 (0.5 mg/mL) and anti-CD49d (0.5 mg/mL) antibodies. Where indicated, cells were stimulated with PMA (Sigma-Aldrich, 50 ng/mL) and ionomycin (Sigma-Aldrich, 1 µg/mL), as positive control. IFN-γ formation was measured by flow cytometry or ELISPOT. For flow cytometry, 1 hour after re-stimulation with peptides, cells were added BD GolgiStop^TM^, containing monensin and BD GolgiPlug^TM^, containing brefeldin A according to manufacturer’s suggestion. IFN-γ production was measured 12-hours after the addition of protein transport inhibitors by intracellular staining usingBD Cytofix/Cytoperm^TM^ reagents according to manufacturer’s protocol. For ELISPOT analysis, cells were re-stimulated in plates with mixed cellular ester membrane that were coated with anti-IFN-γ antibody (Mabtech, 4 µg/mL). Plates were processed for IFN-γ detection after 48-hours of culture.

## Author Contributions

N.B. and S.B. initiated the project; N.B, V.R., C.C.B and T.O. designed the study; C.C.B. collected the samples; V.R and C.C.B. acquired and analyzed data. N.B, B.G., V.R., C.C.B and T.O interpreted the data; V.R. and C.C.B. wrote the manuscript; V.R., C.C.B., S.B., B.G. and N.B. revised the manuscript.

## Supplementary Information

**Supplementary Figure 1.**
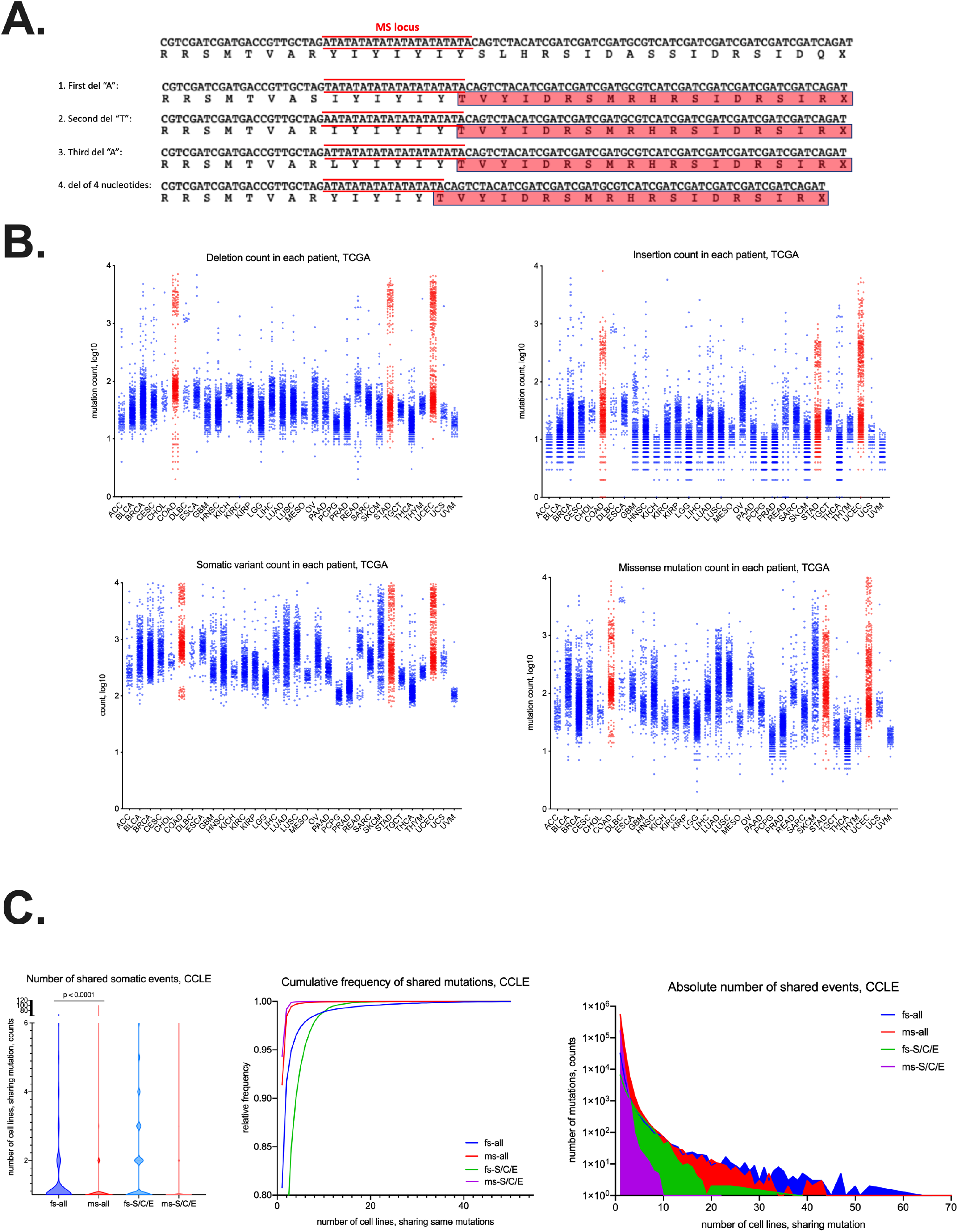
Mutational analysis of TCGA. **A.** Hypothetical example of multiple deletions happening in microsatellite region and leading to identical frameshift (fs-)peptide. **B.** Comparison of frequencies of shared somatic indels and missense point mutations in human cancer cell lines from CCLE. LEFT – number of cancer cell lines, sharing same fs-mutation or same missense point mutation from all CCLE (fs-all and ms-all respectively) or from stomach, colon and endometrium cell lines (fs-S/C/E and ms-S/C/E respectively). Statistical significance derived from non-parametric Mann-Whitney, one-tailed test. CENTER – cumulative frequency plot of shared indel and missense mutation from all CCLE (fs-all and ms-all respectively) or from stomach, colon and endometrium cell lines (fs-S/C/E and ms-S/C/E respectively). RIGHT – absolute number of shared mutations in cancer cell lines from all CCLE (fs-all and ms-all respectively) or from stomach, colon and endometrium cell lines (fs-S/C/E and ms-S/C/E respectively). **C.** Mutational load across different tumor types in TCGA. From left to right, top to bottom: deletion, insertion, somatic variant and missense mutation load.

**Supplementary Figure 2.**
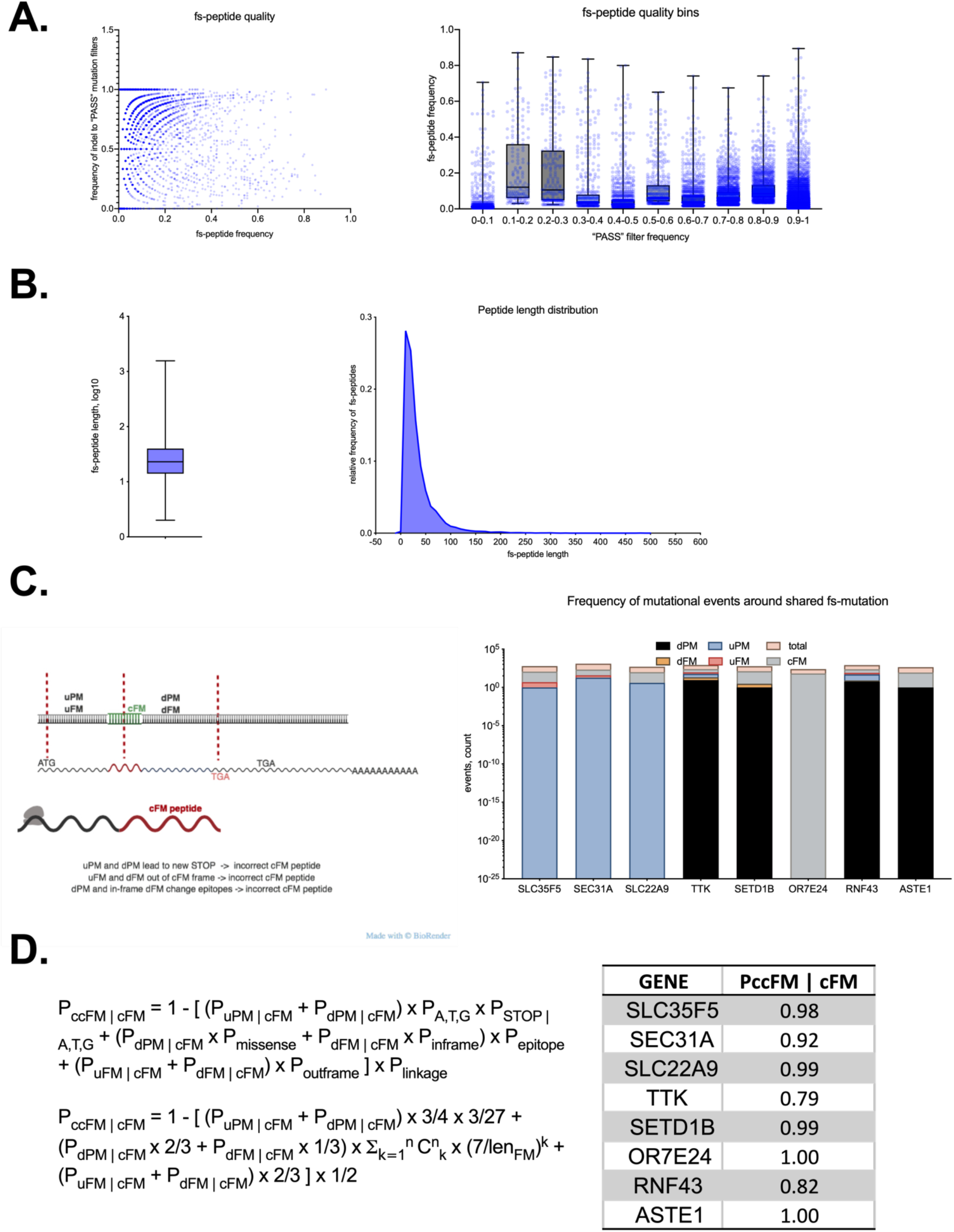
Properties of predicted frameshift peptides. **A.** Distribution of quality metrics for each frameshift mutation detected in MSI-H patients. LEFT – ratio of “PASS” mutation calls for each frameshift, plotted against fs-mutation frequency. RIGHT – box-plot of frameshift frequency in MSI-H patient cohort per 10%-quality bin. The least shared fs-events are the most confident, while majority of shared fs-mutations are within 10-50% “PASS” quality range. **B.** Box plot (LEFT) and distribution (RIGHT) of fs-peptide frequency as function of its length. Most frequent fs-peptides are in 15-50 aminoacid residue range. **C, D.** Estimation of correctness of nine shared frameshift peptides in MSI-H UCEC population. In the reduced approximation, any upstream mutation (upstream fs-mutation uFM or upstream point mutation uPM) may alter the fs-peptide if uFM appears somewhere between start codon and predicted shared fs-mutation or if uPM leads to an abortive stop codon; alternatively, any downstream mutation (downstream fs-mutation dFM or downstream point mutation dPM) may alter shared fs-peptide if dFM happens between shared mutation and novel stop codon, defined by the frame of fs-peptide or if dPM happens within predicted T cell epitope or leads to abortive stop codon. **C.** LEFT: representation of upstream and downstream mutagenesis, which can be detrimental to the predicted fs-peptide; RIGHT – quantification of detrimental events around shared fs-peptides. **D.** LEFT: probabilistic function to estimate conditional probability of fs-peptide being correct given that this fs-mutation is happened; RIGHT: estimated posterior probabilities for each nine shared fs-peptides.

**Supplementary Figure 3.**
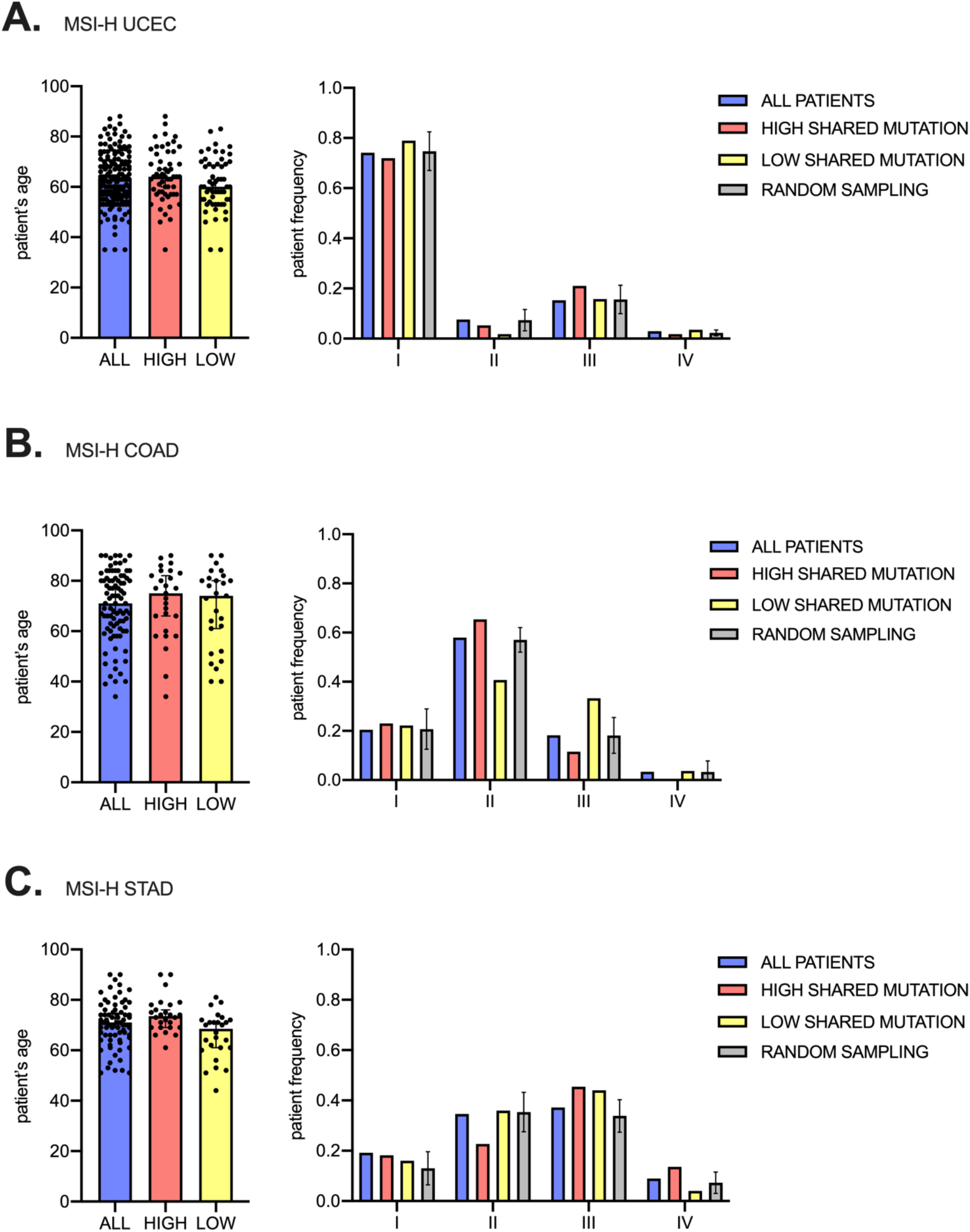
Distribution of patient’s age and tumor stages based on shared fs-epitope load in MSI-H tumors, TCGA. **A.** LEFT – distribution of patient’s age in all, shared fs-epitope high and shared fs-epitope low patients’ cohort, MSI-H UCEC tumor type. RIGHT – patients’ frequency across tumor stages in the same cohorts, as above. **B.** LEFT – distribution of patient’s age in all, shared fs-epitope high and shared fs-epitope low patients’ cohort, MSI-H COAD tumor type. RIGHT – patients’ frequency across tumor stages in the same cohorts, as above. **C.** LEFT – distribution of patient’s age in all, shared fs-epitope high and shared fs-epitope low patients’ cohort, MSI-H STAD tumor type. RIGHT – patients’ frequency across tumor stages in the same cohorts, as above.

**Supplementary Figure 4.**
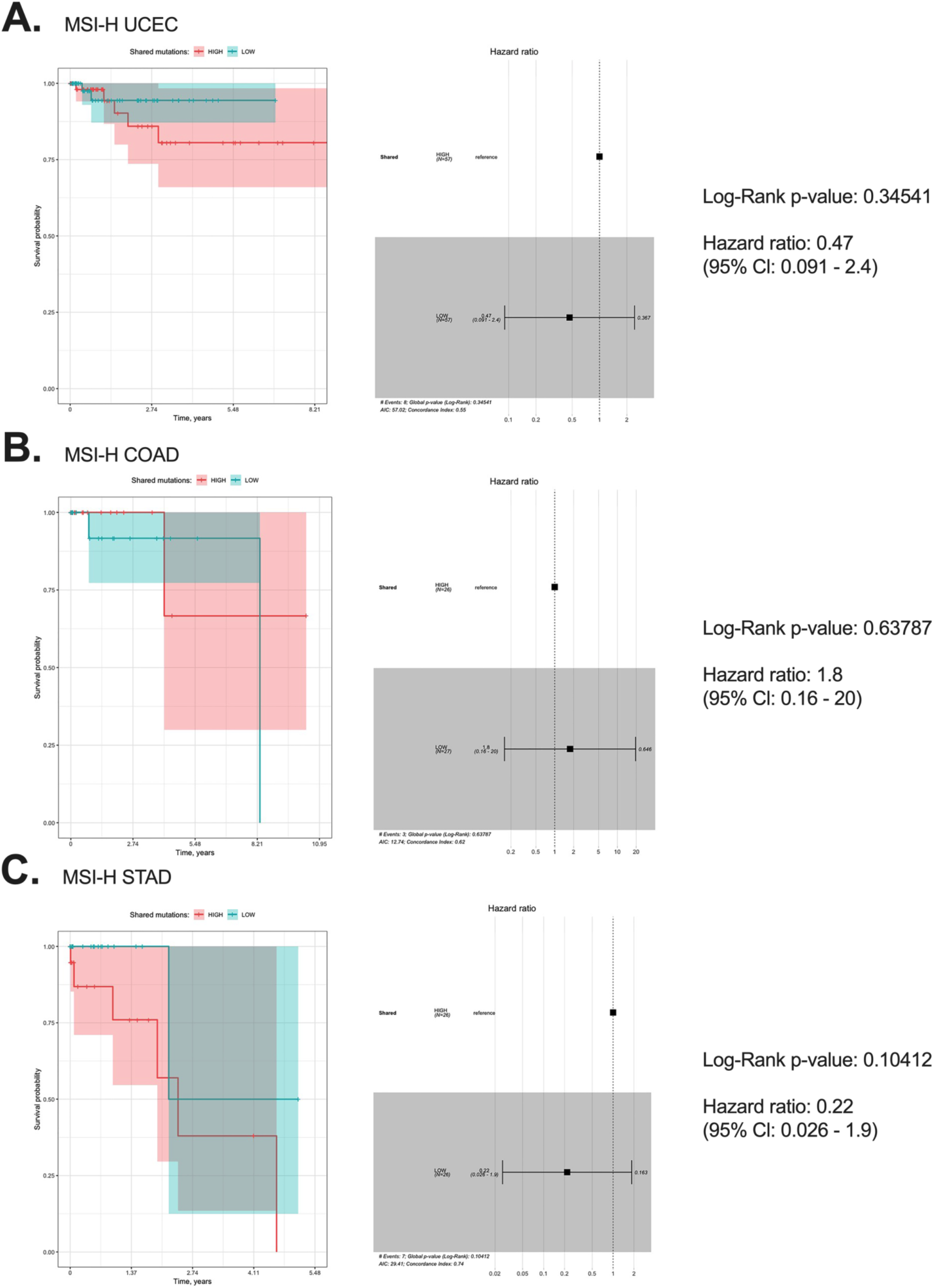
Kaplan-Meier plot of survival analysis (LEFT) and Cox’s proportional hazards model (RIGHT) of MSI-H UCEC (**A**), COAD (**B**) and STAD (**C**) patients, segregated by shared fs-epitope load (HIGH vs LOW).

**Supplementary Figure 5.**
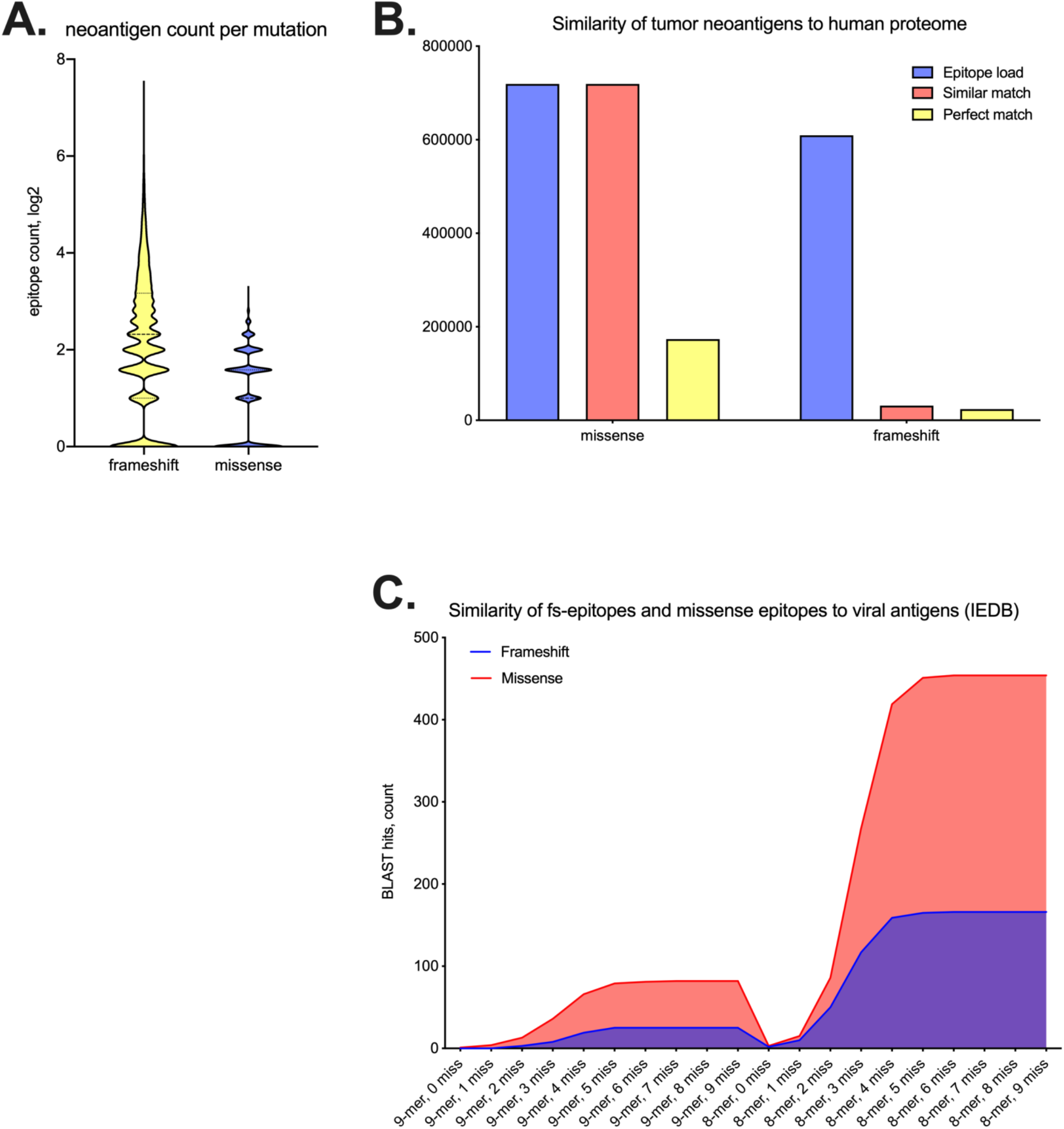
Comparison of missense epitopes and fs-epitopes in MSI-H patients with human proteome and annotated viral epitopes (IEDB). **A.** Quantification of epitopes per missense or frameshift mutation. On average, frameshift mutation generates 4 epitopes, while missense - 2. **B.** Mapping missense and fs-epitopes back to human proteome (ensemble, 2016). “Epitope load” indicates total amount of predicted neoantigens from missense and frameshift mutations; “Similar match” shows number of epitopes, successfully aligned to human proteome by sequence of 8 non-gapped aminoacid residues, allowing 1 mismatch; “Perfect match” shows number of epitopes, “Perfect match” shows number of epitopes, perfectly matched to human proteome by sequence of 9 non-gapped aa with no mismatches. **C.** BLAST comparison of missense epitopes and fs-epitopes with viral epitopes from IEDB, allowing different number of mismatches. Missense-derived epitopes are more similar to viral epitopes then frameshift-derived on average.

**Supplementary Figure 6.**
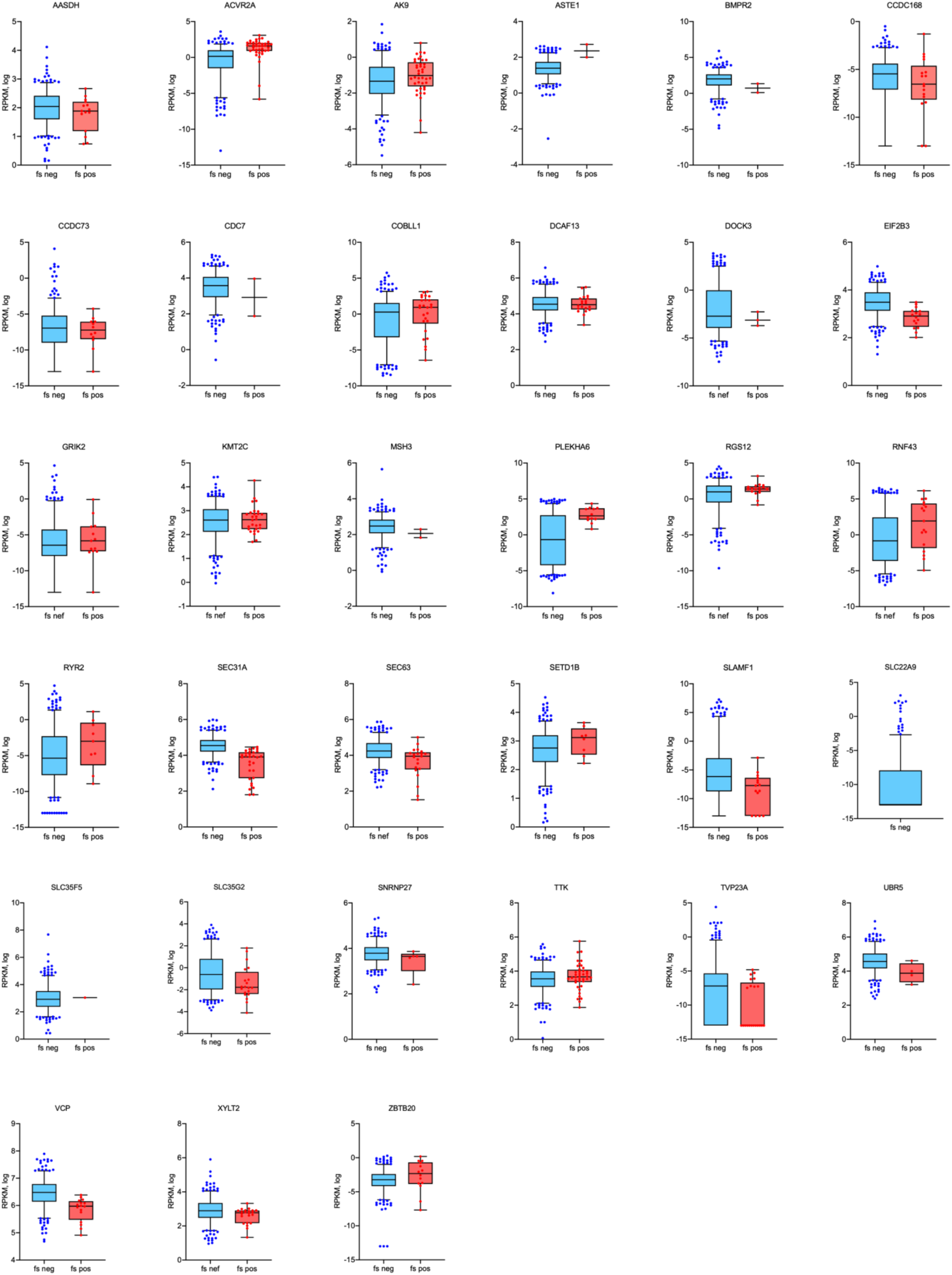
RNA expression (RPKM, log2) of genes, carrying shared fs-mutation in cell lines derived from CCLE. Each box-plot represents RPKM expression of gene of interest in shared fs-negative and shared fs-positive cell lines.

**Supplementary Figure 7.**
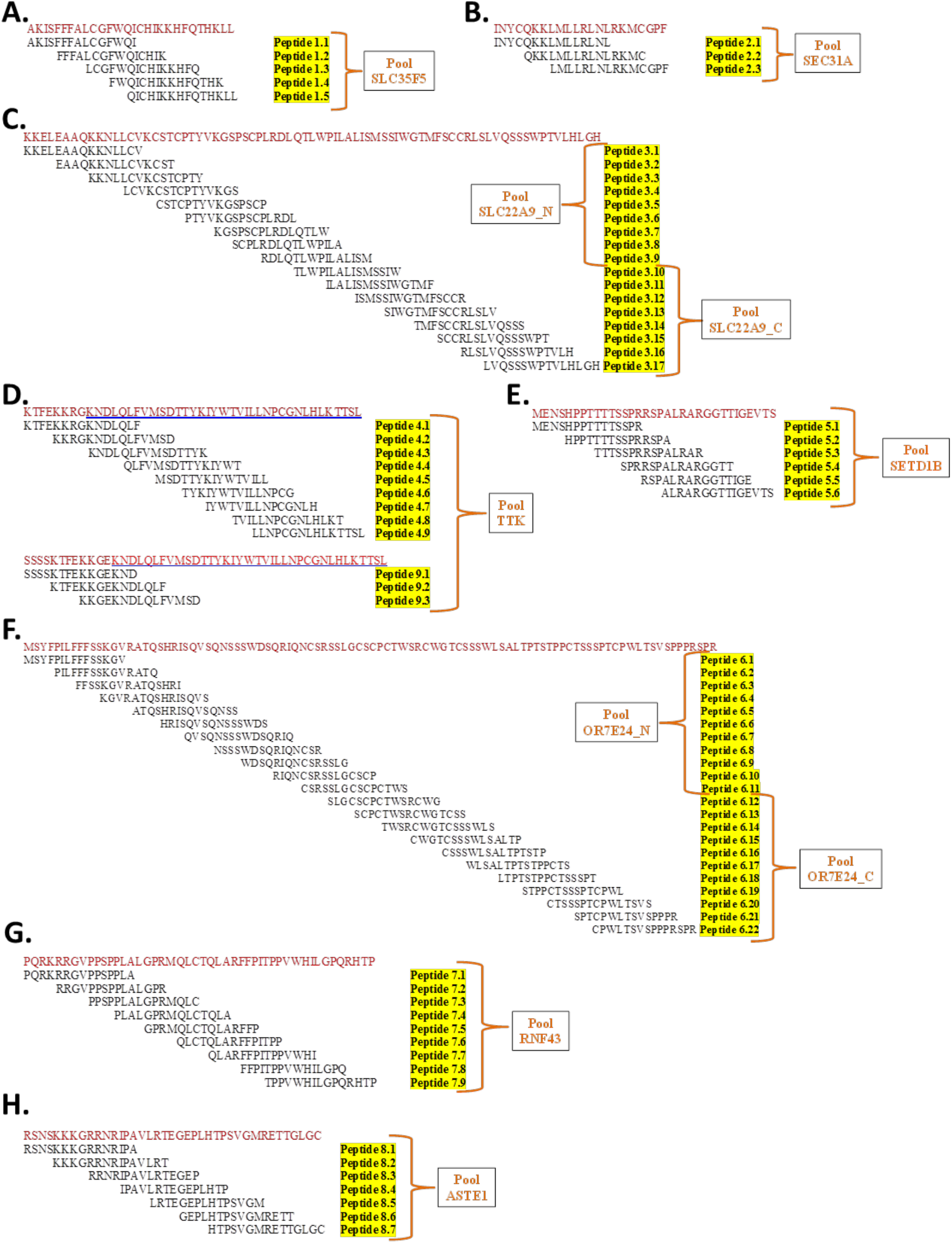
Peptide design. 15-mer overlapping peptides spanning the upstream 8aa WT sequence and the entire mutated fs-peptide were synthesized for the predicted nine shared fs-peptides from MSI-H UCEC. Peptide pools as they were utilized in the immunogenicity assays were denoted. **A.** SLC35F5, **B.** SEC31A, **C.** SLC22A9**, D.** TTK**, E.** SETD1B, **F.** OR7E24, **G.** RNF43, **H.** ASTE1.

**Supplementary Figure 8.**
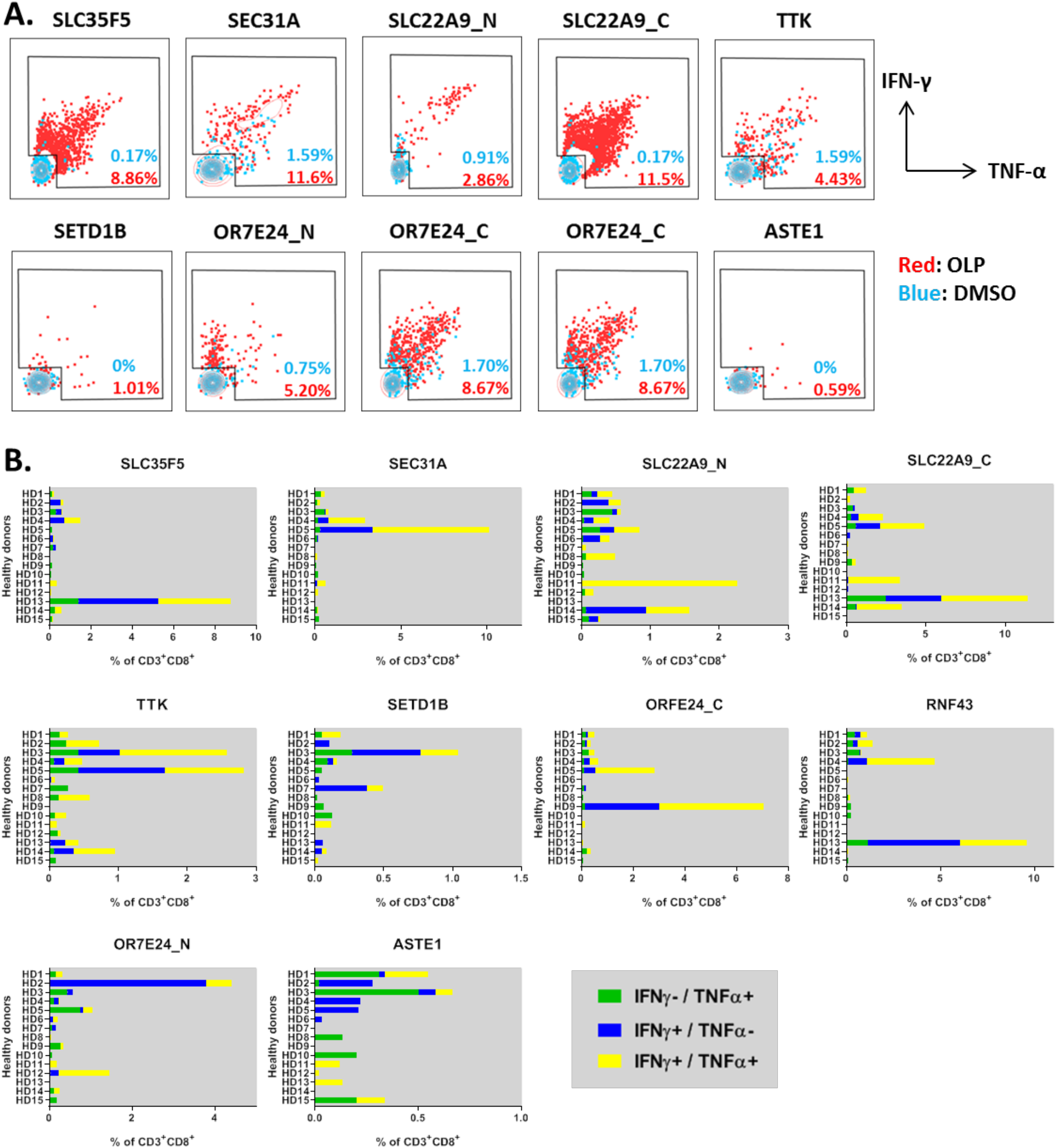
Fs-peptide-specific T cells are polyfunctional. PBMCs from healthy donors (HD) were expanded *in vitro* following stimulation with fs-peptide OLPs. DMSO was used vehicle control. **A.** Representative flow cytometry overlay plots for IFN-γ (y axis) and TNF-α (x axis) production by CD8+ T cells. Cytokine production upon OLP or DMSO stimulation was shown in red or blue, respectively. **B.** Frequencies of CD8+ T cells producing IFN-γ or TNF-α in response to each fs-peptide OLP pool were plotted for each subject. Green denotes HD producing only TNF-α, blue denotes HD producing only IFN-γ and yellow denotes HD producing both IFN-γ and TNF-α. % IFN-γ values were calculated by subtracting the values obtained after DMSO stimulation from the values after OLP pool stimulation and negative values were set to zero.

